# Tumor-specific Causal Inference (TCI): A Bayesian Method for Identifying Causative Genome Alterations within Individual Tumors

**DOI:** 10.1101/225631

**Authors:** Gregory Cooper, Chunhui Cai, Xinghua Lu

## Abstract

Precision medicine for cancer involves identifying and targeting the somatic genome alterations (SGAs) that drive the development of an individual tumor. Much of current efforts at finding driver SGAs have involved identifying the genes that are mutated more frequently than expected among a collection of tumors. When these population-derived driver genes are altered (perhaps in particular ways) in a given tumor, they are posited as driver genes for that tumor. In this technical report, we introduce an alternative approach for identifying causative SGAs, also known as “drivers”, by inferring causal relationships between SGAs and molecular phenotypes at the individual tumor level. Our tumor-specific causal inference (TCI) algorithm uses a Bayesian method to identify the SGAs in a given tumor that have a high probability of regulating transcriptomic changes observed in that specific tumor. Thus, the method is focused on identifying the tumor specific SGAs that are causing expression changes that are specific to the tumor. Those SGAs that have a high probability of regulating transcriptomic changes related to oncogenic processes are then designated to be the putative drivers of the tumor. In this paper, we describe in detail the TCI algorithm and its implementation.

## 1. Introduction

Cancer is mainly caused by SGAs, such as somatic mutations (SMs), somatic copy number alterations (SCNAs), chromosome rearrangement and other genomic alterations. A tumor cell commonly hosts hundreds to over a thousand SGAs, among which only a small minority contribute to tumor development by perturbing cellular signaling pathways while most others are passenger SGAs (unrelated to cancers). A foremost task of precision oncology for cancer treatment is to identify and target the driver SGAs of an individual tumor. Current methods of identifying candidate driver SGAs are mostly based on the assumption that, if a gene is mutated at a frequency significantly above the expected rate in a cohort of tumors, the mutation events of the gene are likely positively selected in tumors due to resultant oncogenic advantages. Therefore, such a gene is more likely a cancer driver gene [1-4]. Hereafter, we refer to this family of methods as *frequency-oriented models*. These models do not attempt to explicitly determine the functional role of a driver in cancer development, that is, they cannot provide insight into functional impact of oncogenic processes caused by a driver SGA. In general, frequency-oriented models are constrained by the need to define the baseline mutation rate, and different models for estimating the baseline rate will lead to different results.

It is well accepted that driver genes can contribute to cancer development through various types of genomic alterations, such as chromosome structure variations, non-coding mutations, and epigenetic modifications [3, 5-7]. For example, copy number amplification and promoter mutations of the telomere reverse transcriptase (*TERT*) play important roles in different cancer types [8, 9]. However, to our knowledge, there is no reported principled method to integrate multiple types of SGA events to determine the significance of the corresponding gene in cancer development, nor there is any theoretical method that can systematically infer the functional impact of driver SGAs perturbing a common gene.

Here, we introduce a novel framework that identifies driver SGAs in a tumor-specific and signal-oriented fashion. Our approach is based on the assumption that driver SGAs cause cancer progression by perturbing signaling pathways, and as such their functional impact is reflected by the cellular or molecular phenotypes regulated by these perturbed pathways. Thus, the task is to find the SGAs that causally regulate cancer-related molecular phenotypes, e.g., differential expression of genes involved in oncogenic processes, for each individual tumor. To this end, we designed a tumor-specific causal inference (TCI) algorithm that infers causal relationships between SGAs and differentially expressed genes (DEGs) within a specific tumor.

The Bayesian causal inference framework developed in this study provides a principled approach to not only incorporate biological prior knowledge and theoretical assumptions but also integrate diverse types of genomic and molecular phenotypic data to infer the functional impact of genomic alterations in individual tumors [10, 11]. In these respects, TCI first calculates the prior probability that an SGA is a driver in the tumor of interest. Based on the positive selection assumption underlying the frequency-based methods, we assume that the more often are the SGA events perturbing the corresponding gene in a tumor cohort, the more likely the gene is a driver in the current tumor. As such, the calculation of the prior incorporates the strength of the frequency-oriented methods [1, 3]. In a signal-oriented fashion, TCI further calculates the marginal likelihood that the molecular phenotype change is caused by the SGA. Finally, TCI derives a posterior probability that the SGA is causally responsible for the observed phenotypic change in a tumor. Thus, TCI unifies the frequency-oriented and signal-oriented approaches to determine the functional impact of an SGA event within a specific tumor.

Previously reported approaches, e.g. eQTL, can infer the association between SGAs and gene expression levels across a population of tumors [12, 13]. To our knowledge, however, no previously published method is capable of inferring the causal relationships between SGAs and differentially expressed genes (DEGs) in a tumor-specific manner. In this paper, we introduce such an approach which is tumor specific in two ways. First, for a given DEG *E* in a given tumor *t*, the only SGAs that can possibly cause (drive) *E* are the SGAs in *t*; SGAs that occur in other tumors, but not in *t*, are not candidate drivers of *E* in *t*. Thus, the search space for candidate drivers is focused in a tumor-specific manner. Second, the scoring of a given SGA in *t* being a driver of *E* is scored probabilistically in manner that is tumor-specific, as we explain in Section 2.2.

The change in going from population-based to tumor-specific causal inference is substantial. Since multiple SGAs can perturb a common signaling pathway, we should consider the causal relationships between SGAs perturbing the pathway and a DEG regulated by the pathway as a multiple-to-one relationship. For example, multiple perturbations of the PI3K pathway are known to regulate its downstream gene expression during tumorigenesis [14]. Interestingly, rarely do SGAs perturb a common pathway in an individual tumor, which is a phenomenon referred to as mutual exclusivity [15-19]. Figure 1 illustrates the mutual exclusive tendency among 3 members of the PI3K pathway, *PIK3CA, PTEN*, and *PIK3R1* [14]. At the tumor population level, any SGA perturbing the PI3K pathway can cause the expression change of a downstream gene, while in an individual tumor, it is more likely that only one SGA causes the expression change. The multiple-to-one relationship and the mutual exclusivity of the SGAs significantly dampens the strength of statistical association between an individual SGA and the DEG of interest when viewed across all tumors (perhaps of a given type). Therefore, a conventional EQTL analysis may fail to find the causal relationship between a low frequency SGA (e.g., *PIK3R1*) that is perturbing a downstream gene via the PI3K pathway, because it cannot adequately account for the gene expression variance caused by other common SGAs that are perturbing the gene in other tumors (e.g., mutation/amplification of *PIK3CA* and mutation/deletion of *PTEN*). Although at the population level it may not be significantly (statistically) associated with a DEG *E* that is downstream of the PI3K pathway, an alteration of *PIK3R1* should nonetheless be the most probable cause of *E* in an individual tumor when both *PIK3CA* and *PTEN* are normal. The TCI algorithm takes advantage of such tumor-specific relationships between SGAs and DEGs in order to locate the SGAs that are most likely driving the DEGs.

**Figure 1.**
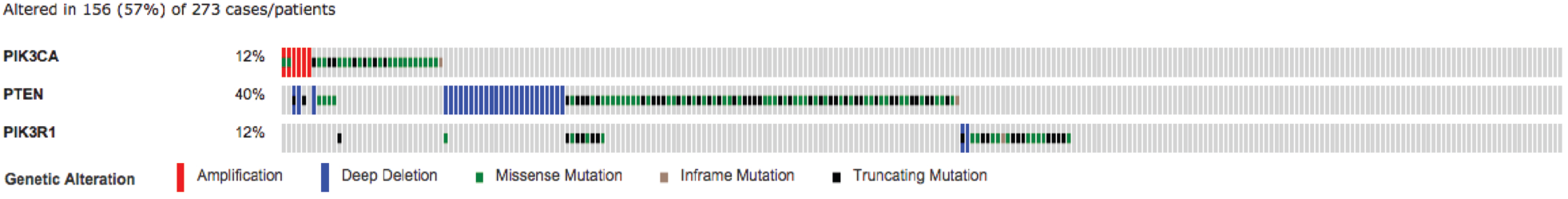
SGAs mutual exclusivity among *PIK3CA*, *PTEN*, and *PIK3R1* in the PI3K pathway. An example of mutual exclusivity among *PIK3CA*, *PTEN*, and *PIK3R1* affecting the PI3K pathway in 273 Glioblastoma Multiforme tumor samples. Each column represents an individual tumor. The combination of color and bar size denotes genetic alteration types: a long red bar represents a tumor with copy number amplification; a long blue bar represents a tumor with copy number deletion; a short green bar represents a tumor with missense mutation; a short brown bar represents a tumor with inframe mutation; a short black bar represents a tumor with truncating mutation. These three SGAs are altered in 156 (57%) out of 273 brain tumors, and as shown, only one of those SGAs occurs in most of the tumors.

## 2. Model Specification

Let the genotypes of all genes in a tumor be represented by a set of binary variables, such that the state of a gene is set to 1 if the gene is altered (e.g., mutated), or otherwise it is set to 0; similarly, let the expression states of all genes be represented by a set of binary variables, such that the expression state of a gene is set to 1 if it is differentially expressed, or otherwise it is set to 0. Let **TS** = {*T*_*1*_, *T*_*2*_, *…, T*_*t*_, *…, T*_*N*_} denote the tumor set which contains a total *N* tumor samples, where *t* indexes over the tumors included in the tumor set. Let **SGA**_*t*_ = {*A*_*1*_, *A*_*2*_, *…, A*_*h*_, *…, A*_*m*_} denote a subset of *m* genes that are altered at the genome level in a tumor *t*, i.e., their genomic states are set to 1, where *h* indexes over the variables in **SGA**_*t*_; let **DEG**_*t*_ = {*E*_*1*_, *E*_*2*_, *…, E*_*i*_, *…, E*_*n*_} denote *n* genes that are differentially expressed in the tumor *t*, where *i* indexes over the variables in **DEG**_*t*_. Hereafter, we will use **SGA** instead of **SGA**_*t*_ and **DEG** instead of **DEG**_*t*_ for simplicity of notation. For each tumor, we further include a variable *A*_*0*_, to collectively represent non-specific factors other than SGAs (e.g., tumor microenvironment) that may affect the gene expression in a tumor. Based on the assumptions that each DEG is likely to be regulated by one aberrant signaling pathway and such a pathway is likely perturbed by only one SGA observed in the tumor (mutual exclusivity), the TCI model further constrains each DEG to be causally regulated by only one SGA (or by *A*_*0*_) in a given tumor. The TCI model assumes no hidden common causes among the variables in **SGA⏝DEG**, including the presence of mixture distributions. It is not concerned with modeling the causal relationships among the variables within **DEG** or among the variables within **SGA**. With the above settings, the task is to determine for each variable *A*_*h*_ in **SGA** the probability that it is the cause of one or more variables in **DEG**, which we interpret as the probability that *A*_*h*_ is a driver in tumor *t*.

For a given tumor, we represent the causal relationships between the variables in **SGA** and those in **DEG** using a bipartite causal Bayesian network (CBN) in which the variables in **SGA** are at level 1 and the variables in **DEG** are at level 2. In such a CBN, arcs always point from **SGA** to **DEG**. A permissible CBN model *M* has only one arc coming into each variable *E*_*i*_ in **DEG** from one variable *A*_*h*_ in **SGA** or *A*_*0*_ which means that it is abnormal due to some non-SGA influence. In model *M*, a given *A*_*h*_ can have zero arcs (a passenger SGA) or one or more arcs emanating from it to the variables in **DEG**; thus, an SGA can causally regulate multiple DEGs. Figure 2 shows an example of a permissible model. Since each tumor generally has a unique **SGA** set and a unique **DEG** set, the model is called tumor-specific.

**Figure 2.**
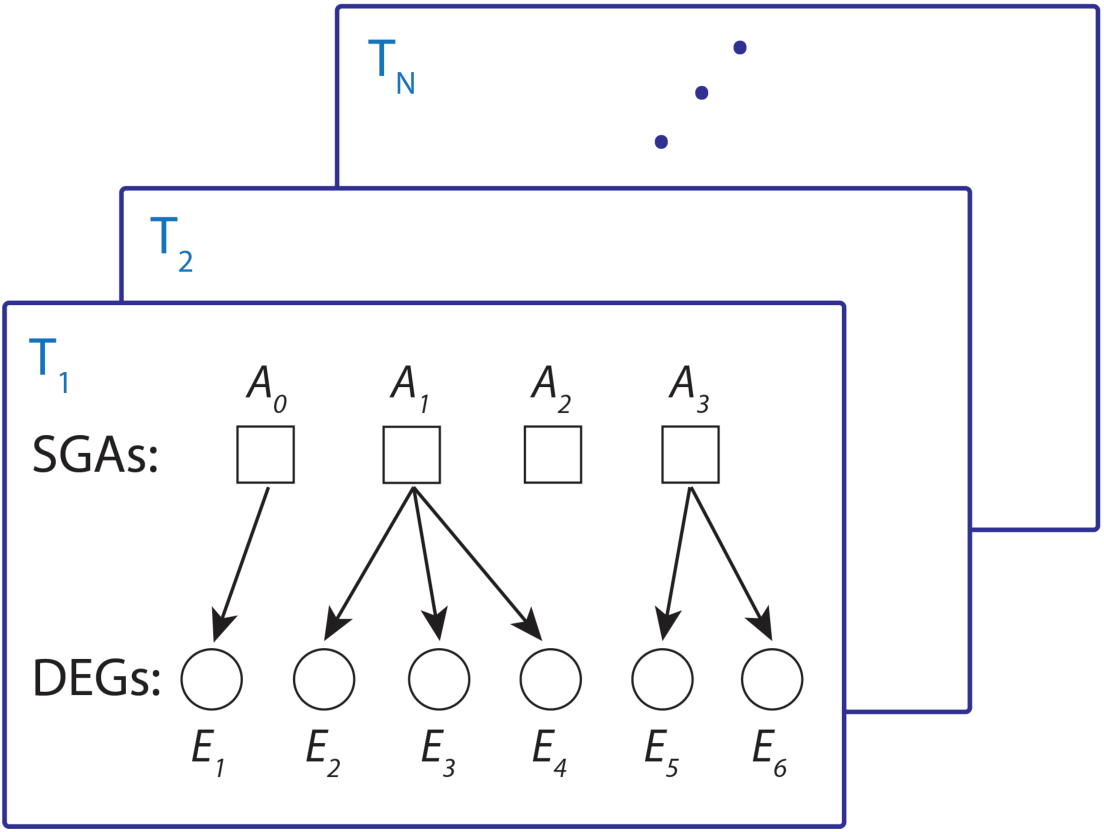
Tumor-specific Bayesian causal inference framework. An example of an admissible CBN for inferring causal relationships between SGAs and DEGs. Here, each plate represents one tumor. For the tumor *T*_*1*_, set **SGA** has three SGA variables plus the non-specific factor *A*_*0*_ (m = 4) and set **DEG** has six DEG variables (n = 6). Each *E*_*i*_ must have exactly one arc into it, which represents having one cause among the variables in set **SGA**. In this model, *E*_*1*_ is caused by *A*_*0*_, and *A*_*1*_ and *A*_*3*_ are drivers of DEGs ({*E*_*2*_, *E*_*3*_, *E*_*4*_} and {*E*_*5*_, *E*_*6*_} respectively), while *A*_*2*_ is not a driver.

### 2.1 The basic Bayesian framework of the TCI model

Let *M* be a CBN structure and let *D* be an observational training dataset, in which each case denotes a sample that contains a measurement for each of the variables in *M*. We assume that the cases in *D* are i.i.d.

We can derive the posterior probability of a CBN structure *M* as follows:

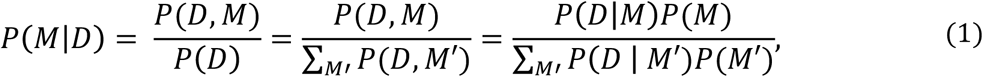

where the sum is taken over all admissible models *M’*. The term *P*(*M*) denotes prior belief that the data-generating CBN has *M* as its structure.

We call the term *P*(*D, M*) the score of CBN structure *M* in light of data *D*. As shown in Equation 1, it can be expressed as follows:

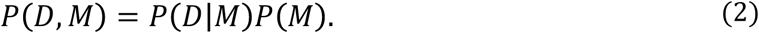

We will assume that *P*(*M*) is a modular prior probability that can be expressed as follows:

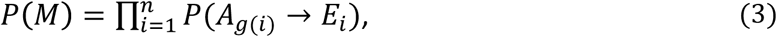

where *A*_*g*(*i*)_ is a node in **SGA** that is the parent of *E*_*i*_ in *M*, and *P(A*_*g*(*i*)_*->E*_*i*_) is the prior probability that *A*_*g*(*i*)_ is causally influencing *E*_*i*_ in the current tumor. The function *g*(*i*) returns an index of *A*. If *g*(*i*) = 0 then *A*_*0*_ represents its parent, which means *E*_*i*_ is regulated by a non-SGA factor in the tumor.

The term *P*(*D*|*M*) is the marginal likelihood of *M*, which can be derived by marginalizing out model parameters *θ* as follows:

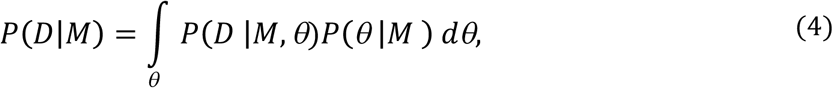

where *θ* represents the parameters (probabilities) associated with CBN structure *M*.

If we assume parameter independence, parameter modularity, and Dirichlet prior probability distributions, we can solve Equation 4 to derive *P*(*D*|*M*) in closed form[20] as follows:

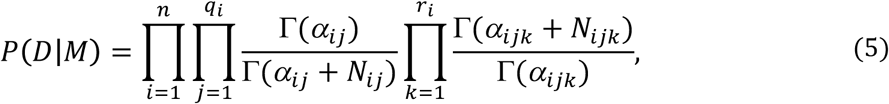

where:

- *i* indexes over the **DEG** variables included in *M*;
- *n* is the number of DEGs in *M*, i.e., the nodes in the **DEG** set of *M*;
- *j* indexes over the 0/1 values (states) of a gene *A* in **SGA** that is being modeled as the parent of *E*_*i*_ in *M*;
- *q*_*i*_ is the number of values of parent gene *A* of the node *E*_*i*_, which is 2, because the *A* is modeled as having the values 1 (a somatic genome alteration) and 0 (no alteration);
- *k* indexes over the 0/1 values of the expression states of *E*_*i*_;
- *r*_*i*_ is the number of values of node *E*_*i*_, which is 2, because *E* is modeled as having the values 1 (a differential gene expression level) and 0 (a normal level of gene expression);
- *N*_*ijk*_ is the number of cases in *D* that node *E*_*i*_ has the value denoted by *k* and its parent has the value denoted by *j*;
- *α*_*ijk*_ is a parameter in a Dirichlet distribution that represents prior belief about *P(E*_*i*_ | *parent(E*_*i*_)); it can be interpreted as belief equivalent to having previously seen (prior to data *D*) *α*_*ijk*_ cases in which node *E*_*i*_ has the value *k* and its parent has the value *j*;
- Γ is the gamma function;
- 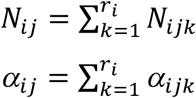
- 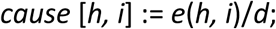

Substituting Equations 3 and 5 into Equation 2 yields:

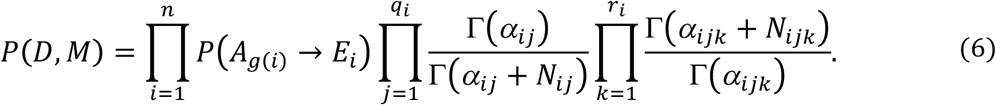

Let *e*(*g*(*i*), *i*) represent the function that appears inside the outer product of Equation 6. Thus, it is defined as follows:

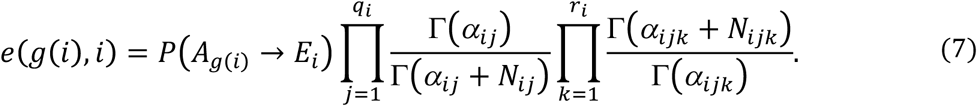

We can now write Equation 6 as follows:

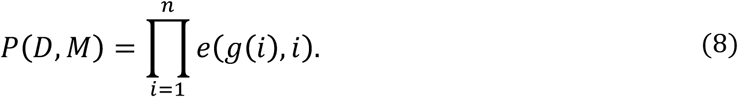

Equation 7 is the score for a causal arc existing from *A*_*g*(*i*)_ to *E*_*i*_. However, we wish to have a non-zero score only for a causal relationship that satisfies the following constraint: *E*_*i*_ is more likely to be abnormal (value 1) when *A*_*g*(*i*)_ is abnormal (value 1) than when *A*_*g(i)*_ is normal (value 0). Given the Dirichlet distributions we are using, the expectation of these conditional probabilities is as follows:

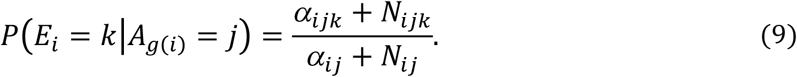

Using conditional probabilities of this form to enforce the constraint mentioned above, leads to the following function:

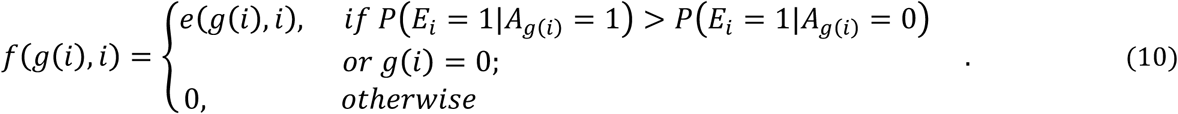

We next use *f* to refine Equation 8 to be the following:

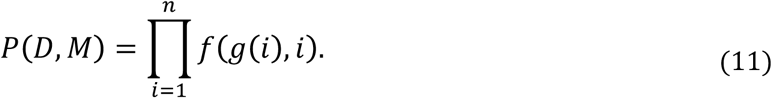

The posterior probability that an SGA *A*_*h*_ causes a DEG *E*_*i*_ in tumor *t* relative to the **SGA** and **DEG** is calculated as follows,

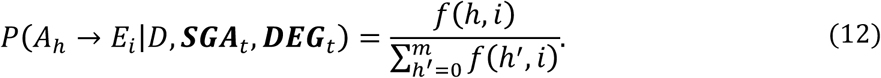

### 2.2 Tumor-centric scoring

When assessing the causal relationship between *A*_*h*_ and *E*_*i*_ using Equation 12, we consider the states of *A*_*h*_ and *E*_*i*_ in all tumors in the training set. As mentioned previously, however, *E*_*i*_ could be regulated by multiple distinct SGAs that affect a common signaling pathway. These SGAs tend to be mutually exclusive across all tumors. For example, a gene that is expressed downstream in the PI3K pathway would be differentially expressed in tumors hosting either a *PTEN* deletion/mutation or a *PIK3CA* amplification/mutations, and these two SGA events tend to be mutually exclusive (Figure 1). Under such circumstances, either a *PTEN* alteration or a *PIK3CA* alteration should be sufficient to explain DEG *E*_*PI3K*_.

In this section, we describe a modified Bayesian scoring measure that models SGAs affecting a DEG. Consider the situation in which *A\** is the driver of DEG *E*_*i*_ in most tumors. Suppose a tumor *t* that is currently being analyzed has *E*_*i*_ as a DEG but does not include *A\** as an SGA. In this case, we need to locate the SGA that is most likely the driver of *E*_*i*_ in tumor *t*, in light of most of the tumors in the training set having *A\** as the driver of *E*_*i*_.

Consider the following example. Let *E*_*PI3K*_ be a DEG in tumor *t*. Suppose the expression of *E*_*PI3K*_ is regulated by the PI3K pathway. Suppose also that *PIK3CA* is the most commonly perturbed member along that pathway (Figure 1), which leads it to be chosen as *A\** according to the methods in Section 2.1. Current tumor *t* does not contain *PIK3CA* as an SGA, however. Thus, we need a causal explanation for DEG *E*_*PI3K*_ in tumor *t*. Suppose that *PIK3R1* is an SGA in tumor *t*. The method described below scores *PIK3R1* as a driver of *E*_*PI3K*_ for all the tumors in the training set that contain *PIK3R1* as an SGA; the remaining tumors in the training set are scored using *PIK3CA* as their driver; the overall score is a function of these two scores. This method is repeated for other SGAs in tumor *t* as candidate causes of *E*_*PI3K*_. If *PIK3R1* turns out to be the SGA in tumor *t* that results in the highest overall score, then *PIK3R1* is output as the most likely driver of *E*_*PI3K*_ in tumor *t*. While this example illustrates the most basic situation in which tumor-specific scoring is called for, the general method can be useful in other circumstances as well.

We now describe the mathematical method we used to implement tumor-specific scoring. In tumor *t*, we want to find the most probable cause of each *E*_*i*_ that has a value of 1 (i.e., is a DEG). Let *A*_*g*(*i*)_ denote a hypothesized gene that is causing *E*_*i*_ to be a DEG in tumor *t*. In order for *A*_*g*(*i*)_ to be a candidate cause, we require that it be altered (i.e., have a value of 1) in tumor *t*. Let 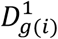 denote the set of tumors in the training set in which variable *A*_*g*(*i*)_ has the value 1, which denotes that these tumors have somatic genome alteration (SGA) in gene *A*_*g*(*i*)_. We can calculate 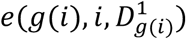 with respect the tumor set 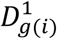 as follows:

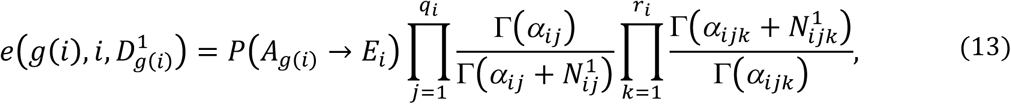

where 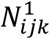 is the number of cases in 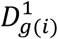 that node *E*_*i*_ has value *k* and its parent *A*_*g*(*i*)_ has value *j*. Since 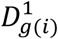 represents the tumors for which *A*_*g*(*i*)_ has the value 1, this means that *j* is fixed at the value 1. Thus, we can simplify Equation 13 to be the following:

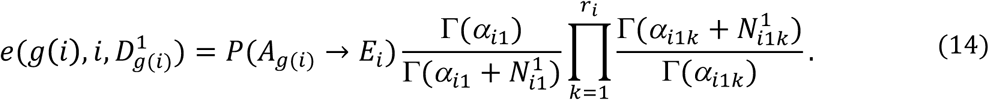

Let 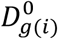 denote the set of tumors in the training set in which variable *A*_*g*(*i*)_ has the value 0, which represents that these tumors do not have genome alteration in *A*_*g*(*i*)_. We need to determine the most likely parent of *E*_*i*_ for these tumors. An efficient way to do so, which we use, is to find the most likely gene *A\** that causes *E*_*i*_ over all tumors in dataset *D*. Then, we hypothesize that gene as the cause of *E*_*i*_ in 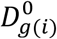 This approach is efficient, because we only have to perform the search once for each *E*_*i*_ prior to seeing the current tumor *t*. More 0(1) specifically, we use the *f* score from Equation 10 (on the data *D* on all the tumors) to search over all possible genes to find the one that maximizes the score. Let *A*_*G*(*i*)_ denote such a maximal scoring gene. We take *A*_*G*(*i*)_ to be the parent of *E*_*i*_ for all the tumors in 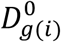 The score for this parent in these tumors in as follows:

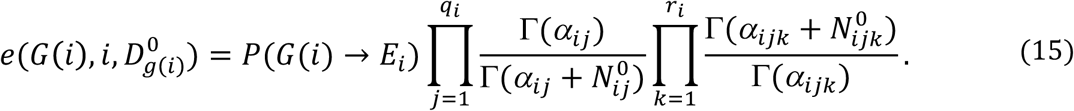

We arrive at a tumor-centric score *e(g(i), i)* by taking the product of Equation 14 and Equation 15:

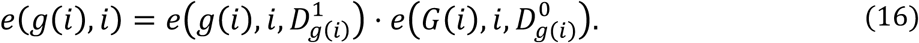

In tumor *t*, consider an *E*_*i*_ that is a DEG (i.e., *E*_*i*_ = 1). We search over all the genes that have a value of 1 (i.e., that are SGAs). For each such *A*_*g*(*i*)_, we use Equation 16 to score it. The *A*_*g*(*i*)_ that has the highest score is returned as the most likely cause of *E*_*i*_ in tumor *t*. We perform this procedure for each *E*_*i*_ that is a DEG in tumor *t*. After doing so, we have determined the most likely SGA causing each DEG in tumor *t*.

## 3. Implementation Details

The following shows the implementation details of our models, i.e., general method and tumor-centric method. Also, in order to apply the methods in the previous section, we need to specify both structure priors and parameter priors.

### 3.1 Pseudocode

The TCI pseudocode in this section consists of a general method and a tumor-centric method. The general method is used to derive the most probable parent *A*_*G*(*i*)_ for each node *E*_*i*_ across all tumors in training set *D*. Then, the tumor-centric method calculates the Bayesian causal score for each edge A_*h*_ ⟶ *E*_*i*_ in each tumor *t*.

#### 3.1.1 General method

~~~
var dataset *D*; set **SGA’**, **DEG’**; array *cause*; integer *i, h, m’, n’*; real *d*;
**SGA’** = the set of genes that have aberrant genome alterations in any tumor of *D*;
*m’* = ∣**SGA’**∣
**DEG’** = the set of genes that are differentially expressed in any tumor of *D*;
*n’* = ∣**DEG’**∣
for *i* = 1 to *n’* do // search for the global driver *A*_*G*(*i*)_ for each *E*_*i*_ at the population level
          for *h* = 0 to *m’* do
                  *compute f*(*h, i*); //use Equation 10 to compute *f*(*h, i*)
          *A*_*G*(*i*)_ is identified as the SGA *G*(*i*) that has the highest *f*(*h,i*) for *E*_*i*_.
~~~

#### 3.1.2 Tumor-centric causal inference

~~~
var dataset *D*; set **SGA**, **DEG**; array *cause*; integer *i, h, m, n*; real *d*;
**SGA** = the set of genes in tumor *t* that have aberrant genome alterations;
*m* = ∣**SGA**∣
**DEG** = the set of genes in tumor *t* that are differentially expressed
*n* = ∣**DEG**∣
for *i* = 1 to *n* do //populate the values of the *cause* array
          *d* = 0,
          for *h* = 0 to *m* do
                  use the global driver *A*_*G*(*i*)_ for *E*_*i*_ as determined using the general method;
                  *d* := *d* + *e*(*h, i*); //use Equation 16 to compute *e*(*h, i*)
          for *h* = 0 to *m* do
                  *cause* [*h, i*] := *e*(*h, i*)/*d*;
~~~

### 3.2 Structure priors

Given the tumor of interest with a unique **SGA** set and a unique **DEG** set, we need to define a tumor specific structure prior *P*(*M*) over permissible CBN structures *M*. Because the **SGA** and **DEG** sets are (with high probability) unique to each tumor, the prior distribution over *M* is also tumor-specific. Assuming the structure prior is modular, we can factorize the *P*(*M*) as a product of prior probabilities for each permissible edge as follows:

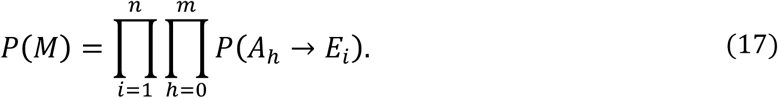

*p*(*M*) comprises a product of prior probabilities of causal edges of a test. A prior probability of a causal edge from a somatic alteration of gene *A*_*h*_ to a DEG *E*_*i*_ can be stated as *P*(*A*_*h*_ ⟶ *E*_*i*_) (abbreviated as θ_*h*_) and determined according to:

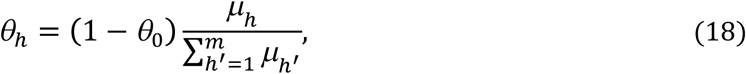

where *θ*_0_ is a prior probability that the cause of DEG *E*_*i*_ is not an SGA, and *h’* indexes over the number *m* of genes in tumor *t* that have SGAs.

Additional genomic information can be applied to derive the prior probability of each edge *A*_*h*_ *⟶ E*_*i*_ using existing prior knowledge. Consider, for example, the availability of the following information for each gene *h*: (1) the number of unique synonymous mutations observed for *h* among the tumors in *D*, and (2) the number of abnormal somatic copy number alterations (according to a given definition of abnormal) of *h* in a normal population without cancer. Such information can be applied to help account for mutation and copy number alterations that are due to differences in gene lengths and chromosome locations. In particular, using the information in (1) and (2) above, we can calculate *μ_h_* as follows:

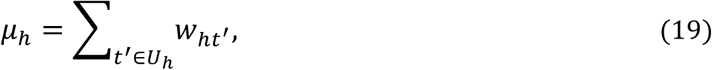

where *U*_*h*_ denotes the tumors in training set *D* that have a somatic alteration in gene *A*_*h*_, and *w*_*ht’*_ denotes a weight proportional to the probability that SGA *h* is a driver in the genome of tumor *t’*. We calculate *w*_*ht’*_ as follows:

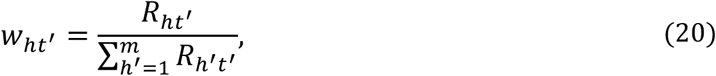

where

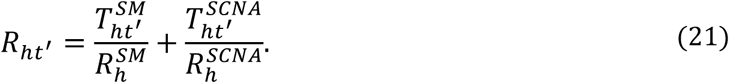

In equation 21, 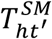 denotes whether gene *h* has a non-synonymous somatic mutation (SM) event or not in tumor *t’*, i.e., 1 or 0, respectively; 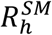 denotes the number of unique synonymous mutation events in gene *h* observed in the reference set of tumor genomes, *D*;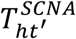 denotes whether gene *h* is affected by an SCNA event or not in tumor *t’*, i.e., 1 or 0, respectively; 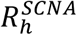 denotes the expected number of times gene *h* is affected by copy number alteration among the tumors in *D*, and yet is only a passenger alteration, based on the number of times gene *h* is affected by copy number alteration in a reference set of cases from a normal human population without known cancer.

### 3.3 Parameter priors

We need parameter priors for when *E*_*i*_ has *A*_*0*_ as its parent and for when it has an *A*_*h*_ in **SGA** set as its parent. Table 1 addresses the case when *E*_*i*_ has *A*_*0*_ as its parent. The Dirichlet parameter values in the table represent that every probability of *P*(*E*_*i*_ *=* 1) is equally likely a *priori* (i.e., before any data are considered); recall that *E*_*i*_ = 1 represents that *E*_*i*_ is abnormal. Table 2 addresses the case in which *E*_*i*_ has some parent *A*_*h*_. The Dirichlet parameter values in the table make it somewhat more likely that *E*_*i*_ will be normal (abnormal) when its cause *A*_*h*_ is normal (abnormal). To make that pattern stronger, the values of 2.0 could be replaced with larger values, such as 2.5, 3.0, or even higher.

**Table 1.** Dirichlet parameter values when *A*_*0*_ is the parent of node *E*_*i*_.

| Prior probability being specified | Dirichlet parameter | Default value |
| --- | --- | --- |
| $P(E_i)$ | $\alpha_{i10}$ | 1.0 |
| | $\alpha_{i11}$ | 1.0 |

**Table 2.** **Dirichlet parameter values when there is one *A*_*h*_ parent of node *E*_*i*_.**

| Prior probability being specified | Dirichlet parameter | Default value |
| --- | --- | --- |
| $P(E_i \mid A_h = 0)$ | $\alpha_{i00}$ | 2.0 |
| | $\alpha_{i01}$ | 1.0 |
| $P(E_i \mid A_h = 1)$ | $\alpha_{i10}$ | 1.0 |
| | $\alpha_{i11}$ | 2.0 |

The computations in this paper have used standard arithmetic operations. However, model scores can become extremely small. Therefore, it is generally necessary to use log arithmetic. When doing so, Equation 7, for example, becomes a sum of log terms, rather than a product of terms.

## Acknowledgement

Research reported in this publication was supported by grant U54HG008540 awarded by the National Human Genome Research Institute through funds provided by the trans-NIH Big Data to Knowledge (BD2K) initiative. The content is solely the responsibility of the authors and does not necessarily represent the official views of the National Institutes of Health.

